# Muscarinic Lateral Excitation Contributes to Visual Object Segmentation during Collision Avoidance

**DOI:** 10.1101/216945

**Authors:** Ying Zhu, Richard B. Dewell, Hongxia Wang, Fabrizio Gabbiani

## Abstract

Visual neurons specialized in tracking objects on a collision course are often finely tuned to their target stimuli as this is critical for survival. The presynaptic neural networks converging on these neurons and their role in tuning them remains poorly understood. We took advantage of well-known characteristics of one such neuron to investigate the properties of its presynaptic input network. We find a structure more complex than hitherto realized. In addition to dynamic lateral inhibition used to filter out background motion, presynaptic circuits include normalizing inhibition and short-range lateral excitatory interactions mediated by muscarinic acetylcholine receptors. These interactions preferentially boost responses to coherently expanding visual stimuli generated by colliding objects, as opposed to spatially incoherent controls, helping implement object segmentation. Hence, in addition to active dendritic conductances within collision detecting neurons, multiple layers of both inhibitory and excitatory presynaptic connections are needed to finely tune neural circuits for collision detection.

## Introduction

Both in vertebrates and invertebrates, collision-detecting neurons are typically located in higher order nuclei and neuropils, several synapses away from the sensory periphery. In fish, for example, collision detecting neurons are found in the optic tectum and the hindbrain (Preuss et al., 2006; Dunn et al., 2016; Temizer et al., 2015). Similarly, in birds they were identified in the nucleus rotundus of the thalamus, a major recipient of ascending tectal projections (Wang and Frost, 1992; Sun and Frost, 1998). In mice and cats, they have been documented in the superior colliculus (Liu et al. 2011; Shang et al. 2015; Zhao et al., 2014). In insects and crustaceans, collision-detecting neurons are typically found in the lobula or the lobula plate of the optic lobes (de Vries and Clandinin, 2012; Wu et al. 2016; Gabbiani et al., 2002; Olivia and Tomsic, 2014), which are considered higher order visual processing neuropils, and in the brain proper, where they convey information to motor centers involved in the generation of escape responses (von Reyn et al. 2014; Rind and Simmons, 1992; Hatsopoulos et al., 1995; Schlotterer, 1977). These arrangements leave plenty of room for presynaptic networks to implement through specific patterns of connectivity the neural computations required to shape the responses of collision-detecting neurons. Although we know much about the response properties of collision-detecting neurons in the systems mentioned above, at present the connectivity patterns of their presynaptic networks and their computational roles remain largely unknown. We took advantage of a well-characterized circuit dedicated to collision avoidance in locusts to investigate these issues.

The lobula giant movement detector (LGMD) in the locust is an identified visual neuron in the third neuropil of the optic lobe of orthopteran insects (O’Shea and Williams 1974). It responds preferentially to objects approaching on a collision course with the animal or their two-dimensional simulations on a screen (i.e., looming stimuli; Hatsopoulos et al. 1995; Rind and Simmons 1992; Schlotterer 1977), and plays an important role in visually evoked escape behavior (Fotowat and Gabbiani 2011; Santer et al. 2006). The LGMD is exquisitely tuned to detect approaching objects on a collision course (e.g., Judge and Rind, 1997; Gray et al. 2001; Gabbiani et al. 2001); a tuning that is in part mediated by active conductances close to its spike initiation zone and within its dendritic tree (Peron and Gabbiani 2009; Dewell and Gabbiani 2017). Since the computation carried out by the LGMD as it tracks approaching objects was first characterized (Hatsopoulos et al. 1995; Gabbiani et al. 1999), a host of neurons with response characteristics resembling those of the LGMD have been described in a variety of species ranging from invertebrate to vertebrate (Sun and Frost 1998; Preuss et al. 2006; Liu et al. 2011; de Vries and Clandinin 2012; Olivia and Tomsic 2014; Temizer et al. 2015; Dunn et al. 2016; von Reyn et al. 2017). In none of these systems, however, do we understand how spatio-temporal interactions within presynaptic circuits shape their responses.

The LGMD neuron receives onto its extended excitatory dendritic field a retinotopic projection of synaptic inputs from an entire visual hemifield. This projection originates from individual ommatidia (facets) on the eye and is mediated by calcium-permeable nicotinic acetylcholine receptors (Peron et al. 2009; Zhu and Gabbiani 2016). Electron microscopy has revealed that the afferents building this excitatory projection (also called trans-medullary-afferents or TmAs) make reciprocal synapses on neighboring afferents at the level of their presynaptic terminals on the LGMD (Rind and Simmons 1998; Rind et al. 2016). Further, histochemical and immunocytochemical methods demonstrated that these lateral connections are cholinergic and suggested that they might mediate inhibition through muscarinic acetylcholine receptors (mAChRs; Rind and Simmons, 1998; Rind and Leitinger, 2000). However, mAChRs can either be excitatory or inhibitory (Brown 2010) and there is to date no direct experimental evidence to support either of these alternatives for the lateral connections presynaptic to the LGMD. Further, there is currently little understanding of the role mAChRs may play in motion detection (Murphy and Sillito, 1991; Soma et al., 2012; Sato et al., 1987; Schmidt, 1992; Shinomiya et al., 2014). This conspicuous pattern of connectivity is intriguing and its computational role remains to be elucidated. One hypothesis is that if the lateral muscarinic connections are inhibitory, the activation of one afferent would inhibit adjacent afferents, setting up a race between excitation and inhibition that might lead to more specific responses for expanding over translating stimuli (Rind and Simmons 1998; Rind et al. 2016). By contrast, if the connections were excitatory, the activation of one afferent would increase synaptic release from adjacent afferents, enabling stronger and longer-lasting responses to coherently expanding stimuli, such as objects approaching on a collision course.

Calcium enters the excitatory dendritic field of the LGMD exclusively through the calcium-permeable nicotinic receptors associated with its afferents (Peron et al. 2009). Thus, in this system calcium imaging offers an accurate means of monitoring activation of the afferents and their lateral interactions. To study the spatiotemporal calcium activation dynamics on the LGMD excitatory dendrites, we used the indicator Oregon Green BAPTA-5N (OGB-5N), whose fast kinetics allowed us to minimize the difference between calcium fluorescence and membrane potential changes. To investigate whether the lateral interactions between TmAs are excitatory or inhibitory, we locally applied the mAChR antagonist scopolamine or the agonist muscarine and tested responses to different visual stimuli. For each stimulus type the lateral interactions proved excitatory. The excitatory nature of lateral connections raises a conundrum: since they amplify an excitatory input that already grows rapidly over the course of object approach, how is this input maintained within the dynamic range of the LGMD? One possibility is for the presynaptic network to rely on global, normalizing inhibition, a feature present in many sensory circuits (e.g., Heeger 1992; Olsen et al. 2010; Rabinowitz et al. 2011). We present several lines of evidence suggesting that such inhibition is indeed part of the connectivity scheme among presynaptic LGMD excitatory afferents. What role does lateral excitation play in collision detection? In a final set of experiments, we show that it helps implement visual object segmentation by tuning the LGMD’s responses to solid objects approaching on a collision course as opposed to spatially incoherent matched controls, a computation critical to elicit appropriate escape behaviors (Dewell and Gabbiani, 2017).

## Results

### Orderly spatio-temporal activation of LGMD excitatory dendritic field is evoked by looming stimuli

The LGMD’s excitatory dendritic field is retinotopically organized (Peron et al. 2009; Zhu and Gabbiani, 2016). We therefore started by mapping the dendritic activation pattern elicited by different regions of the display used to generate visual stimuli (Methods). For this purpose, we presented square flashes at 5 distinct locations (Fig. 1A, top; Supp. Fig. 1A) and determined the dendritic branches activated by each stimulus (Fig. 1A, bottom). Previous work showed that a single dendritic branch is activated by more than one ommatidium (facet) and that the amount of overlap in dendritic activation decreases with inter-ommatidial distance, becoming close to zero for a separation of four ommatidia (Zhu and Gabbiani 2016, Fig. 3). Since each ommatidium is receptive to approximately 2° of visual space when light adapted (Wilson, 1975), we separated the stimuli by 8° to minimize dendritic overlap. Indeed, we found that the branches activated by these stimuli intersected only minimally, as illustrated in Fig. 1A, bottom. How does the expanding shadow pattern of a looming stimulus activate such a dendritic retinotopic mapping? To address this question, we presented looming stimuli with a relatively high half-size (*l*) to speed *(|v|*) ratio, *l/|v|*=120 ms, and thus a long time of approach (Fotowat and Gabbiani, 2007), maximizing our ability to resolve temporally the associated calcium signals (Fig. 1B, top; Supp. Movie 1). These stimuli expanded symmetrically from the center of square 3, as illustrated in Fig. 1A, middle. During stimulus presentation, we simultaneously recorded the calcium responses of the dendrites. The middle panel in Fig. 1B shows for one animal the average calcium responses at the center branches (color-coded red in Fig. 1A, bottom), and two sets of branches symmetrically surrounding them (blue and green Fig. 1A, bottom). The same data is presented on an expanded time scale in Fig. 1C (middle panel).

**Fig. 1.**
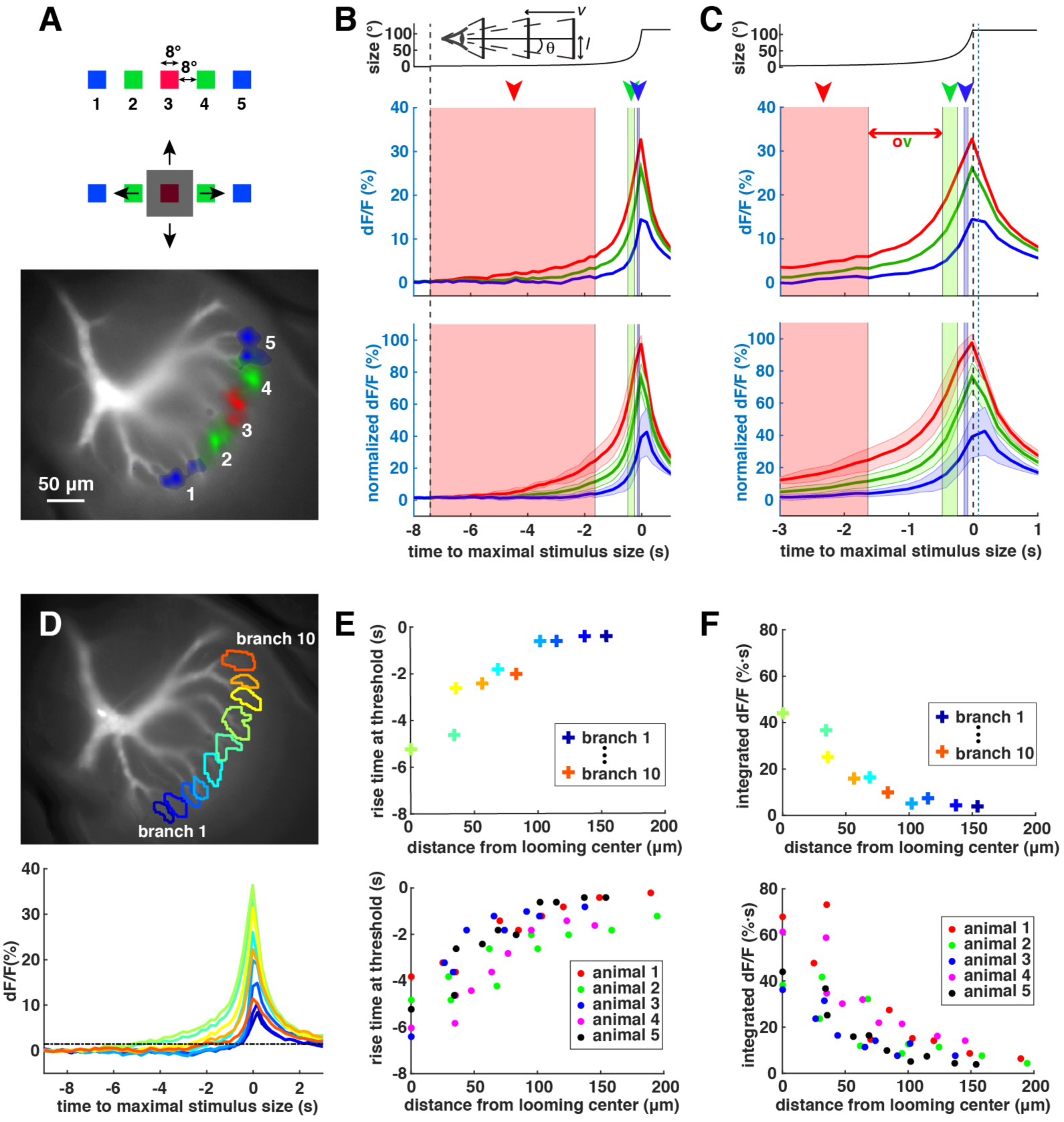
Spatio-temporal activation of the LGMD excitatory inputs in response to looming stimuli. (A)**Top**: to map dendritic receptive fields, brief square flashes were presented at 5 locations individually on the display. Three colors represent flashes with different distances from the display center.**Middle**: a looming stimulus with *l/|v|* = 120 ms was then presented. It expanded from the center square location and eventually covered locations where the other flashes were presented, as illustrated by the schematics. **Bottom**: Excitatory dendritic field of the LGMD. Dendritic branches activated in response to each of the 5 square flashes are shown in red, green and blue colors. (B)The black dashed line indicates the start of the looming stimulus. **Top**: time course of the angular size of the looming stimulus with *l/|v|* = 120 ms. The schematics illustrates the stimulus parameters: radius l, speed v and angular size 2θ. **Middle**: time courses of the average relative fluorescence change dF/F at the center branch (red), and two sets of branches symmetrically surrounding the center branch (green and blue) in response to the looming stimulus in one animal. Time is computed relative to the maximal stimulus size, which occurs 80 ms before projected collision. **Bottom**: mean (solid lines) and standard deviation (shaded areas around the solid lines, n=5 animals) of the time course of the average dF/F (normalized to the maximum dF/F in response to the center square flash for each animal) at the center branch (red), and two sets of branches symmetrically surrounding the center branch (green and blue) in response to the looming stimuli. **Middle and Bottom**: the red, green and blue rectangles (indicated by the red, green and blue arrowheads, respectively) represent the time during which the edges of the looming stimulus moved across the locations of the red, green and blue square flashes, respectively. Note that during the white regions in between the shading, the stimulus edge would be within the receptive fields of branches on both sides. (C)The same as (B) with a zoomed-in time duration. The black dashed line indicates time at which the looming stimulus reached its maximum size, and the dashed blue line the projected time of collision. Red double arrow indicates the time interval during which the edge of the looming stimulus activates both the red and green branches in (A), bottom due to the overlapped activation pattern described in Zhu and Gabbiani (2016). (D)**Top**: excitatory dendritic field of the LGMD with 10 selected dendritic branches indicated by different colors.**Bottom**: time courses of the average dF/F at each of the selected dendritic branches in the **top** with matched colors. The horizontal black dashed line represented a threshold for branch activation which is taken as 5 times the baseline noise level. (E)**Top**: the vertical axis is the time when the rising phase of each trace in (D, **bottom**) crosses the threshold. The branch with the earliest rise time at threshold is defined as the looming center. The horizontal axis is the linear distance in the imaging plane from each selected dendritic branch to the looming center. Colors are matched with the colors of the selected dendritic branches in (D). **Bottom**: same as in **top** but across 5 animals. All the selected dendritic branches are plotted in the same color for each animal. (F)**Top**: integrated dF/F over time for each selected dendritic branch in (D) as a function of distance from the looming center. ***Bottom***: same as in ***top*** but across 5 animals. Data from each animal is plotted in a different color.

**Fig. 3.**
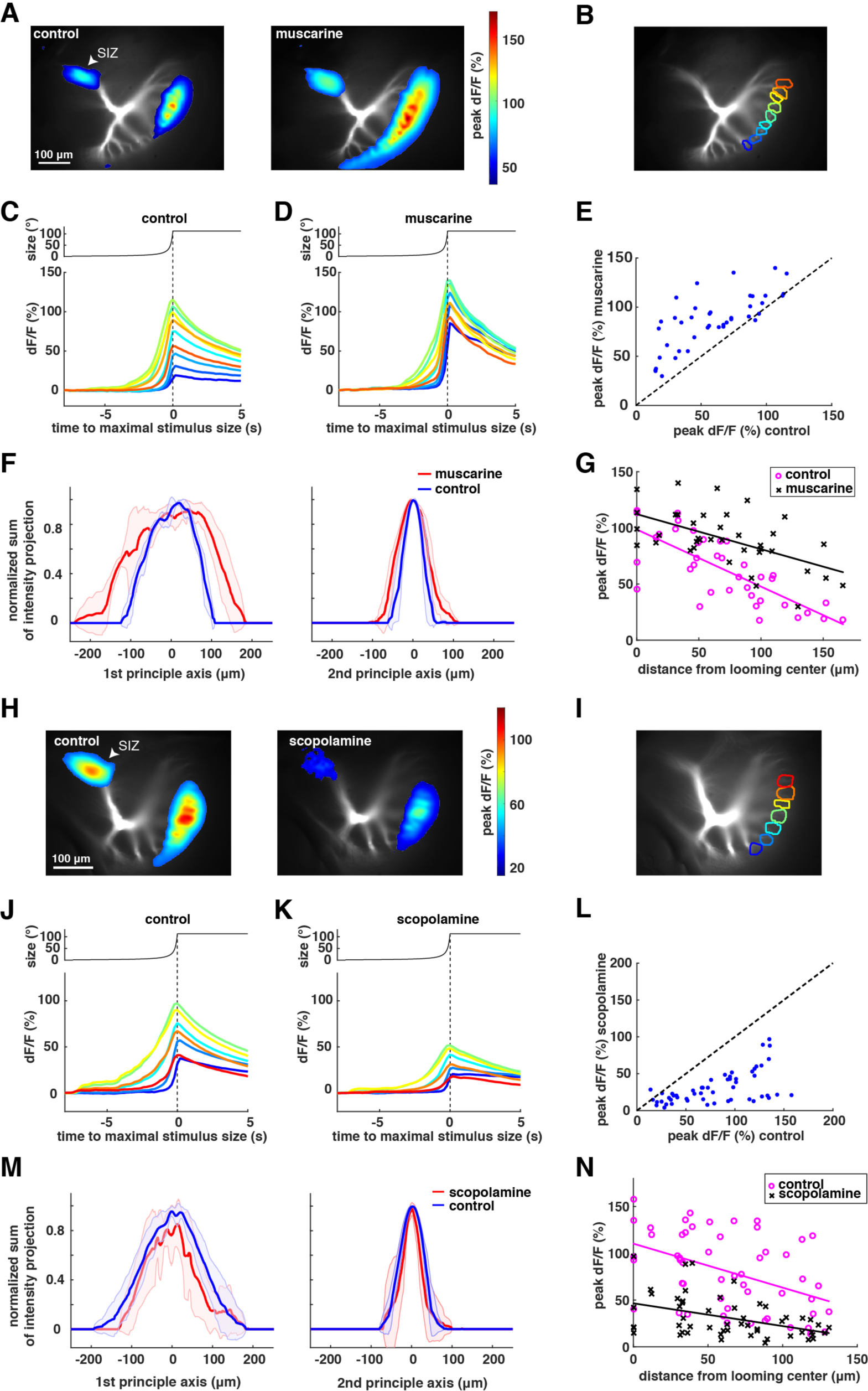
Effect of muscarine and scopolamine on looming responses. (A)***Left***: activation of the LGMD in response to looming. The peak dF/F at each pixel of the activated regions is shown according to the colormap on the right (average over 3 trials in 1 animal). ***Right***: activation of the LGMD in response to looming after puffing muscarine (average over 3 trials, same animal as on left). (B)Excitatory dendritic field of the LGMD with selected dendritic branches indicated by different colors. (C)***Top***: time course of the angular size of the looming stimulus with l/|v|=120 ms. ***Bottom***: time course of the average dF/F in response to looming stimuli at each of the selected dendritic branches in (C), with matched colors in one animal. In (C) and (D), the vertical dashed line indicates the time of maximal stimulus size. (D)Same analysis as in (D) after puffing muscarine. (E)The peak amplitude of dF/F on each selected dendritic branch after puffing muscarine versus before puffing across 5 animals. (p=3.2**⋅**10^−10^, Wilcoxon rank-sum test) (F)***Left***: the sum of the peak dF/F projected onto the first principal axis is plotted as a function of the projection location along that principal axis after normalizing to the peak of the curve without (blue) and with muscarine (red). Red and blue shadings, standard deviation of 5 animals. The FWHM of the curve before muscarine application has a mean of 148.7 µm with a standard deviation of 13.3 µm. The FWHM of the curve after muscarine application has a mean of 252.2 µm with a standard deviation of 13.2 µm. (p=0.0002, paired t-test, before and after muscarine application) ***Right***: same analysis as on the ***Left*** but along the second principal axis. The FWHM of the curve before muscarine application has a mean of 61.2 µm with a standard deviation of 15.7 µm. The FWHM of the curve after muscarine application has a mean of 90.6 µm with a standard deviation of 18.2 µm. (p=0.0016, paired t-test, before and after muscarine application). (G)The peak amplitude of dF/F on each selected dendritic branch is plotted as a function of the distance of that branch to the looming center across 5 animals. Magenta circles, before puffing muscarine (control). Black crosses, after puffing mucarine. Solid magenta and black lines are the linear regression of the data before and after puffing muscarine, respectively. The slopes of the two regression lines are significantly different. (n=38 data points for 5 animals, p=0.023, ANCOVA) (H)***Left***: activation of the LGMD in response to looming. The peak dF/F at each pixel of the activated regions is shown according to the colormap on the right (average over 3 trials in 1 animal). ***Right***: the activation of the LGMD in response to looming after puffing scopolamine (average over 3 trials, same animal as on left). (I)Excitatory dendritic field of the LGMD with selected dendritic branches indicated by different colors. (J)***Top***: time course of the angular size of the looming stimulus with *l/|v|* = 120 ms. ***Bottom***: time course of the average dF/F at each of the selected dendritic branches in response to looming stimuli, with matched colors to (C). (K)Same analysis as in (D) after puffing scopolamine. In (C) and (D), the vertical dashed line indicates the time of maximal stimulus size. (L)The peak amplitude of dF/F on each selected dendritic branch after puffing scopolamine versus before puffing across 5 animals. (p=2.8**⋅**10^−15^, paired t-test) (M) ***Left***: the sum of the peak dF/F projected onto the first principal axis is plotted as a function of the projection location along that principal axis after normalizing to the peak of the curve without (blue) and with scopolamine (red). Red and blue shadings, standard deviation of 7 animals. The FWHM of the curve before scopolamine application has a mean of 168.3 µm with a standard deviation of 39.8 µm. The FWHM of the curve after scopolamine application has a mean of 141.2 µm with a standard deviation of 62.9 µm (p=0.21, paired t-test, before and after scopolamine application). ***Right***: the same analysis as on the ***Left*** but along the second principal axis. The FWHM of the curve before scopolamine application has a mean of 72.9 µm with a standard deviation of 13.1 µm. The FWHM of the curve after scopolamine application has a mean of 60.8 µm with a standard deviation of 31.2 µm. (p=0.36, paired t-test, before and after scopolamine application). (N) The peak amplitude of dF/F on each selected dendritic branch is plotted as a function of the distance of that branch to the looming center across 7 animals. Magenta circles, before puffing scopolamine (control). Black crosses, after puffing scopolamine. Solid magenta and black lines are the linear regression of the data before and after puffing scopolamine, respectively. The slopes of the two regression lines are not significantly different. (n=53 data points for 7 animals, p=0.22, ANCOVA)

Unexpectedly given the retinotopic organization of the excitatory dendritic field, we found that the relative fluorescence change, dF/F, of the center branches (red line in Fig. 1B, middle) reached its peak well after the edge of the looming stimulus had moved outside the borders of the square that they mapped to (red arrowhead and shaded region in Fig. 1B, middle). The calcium signal continued to increase even after the edges of the stimulus had expanded beyond the four lateral squares (blue and green arrowheads and shaded regions in Fig. 1B, C). Furthermore, we found that the central branches that activated earlier also reached a higher peak dF/F. These results were consistent across animals, as illustrated in the bottom panels of Fig. 1B, C. Finally, the peak times of the center and the two sets of surrounding branches were staggered in time (Supp. Fig. 1B), a feature better resolved in the set of experiments we will describe next (Fig. 2J).

**Fig. 2.**
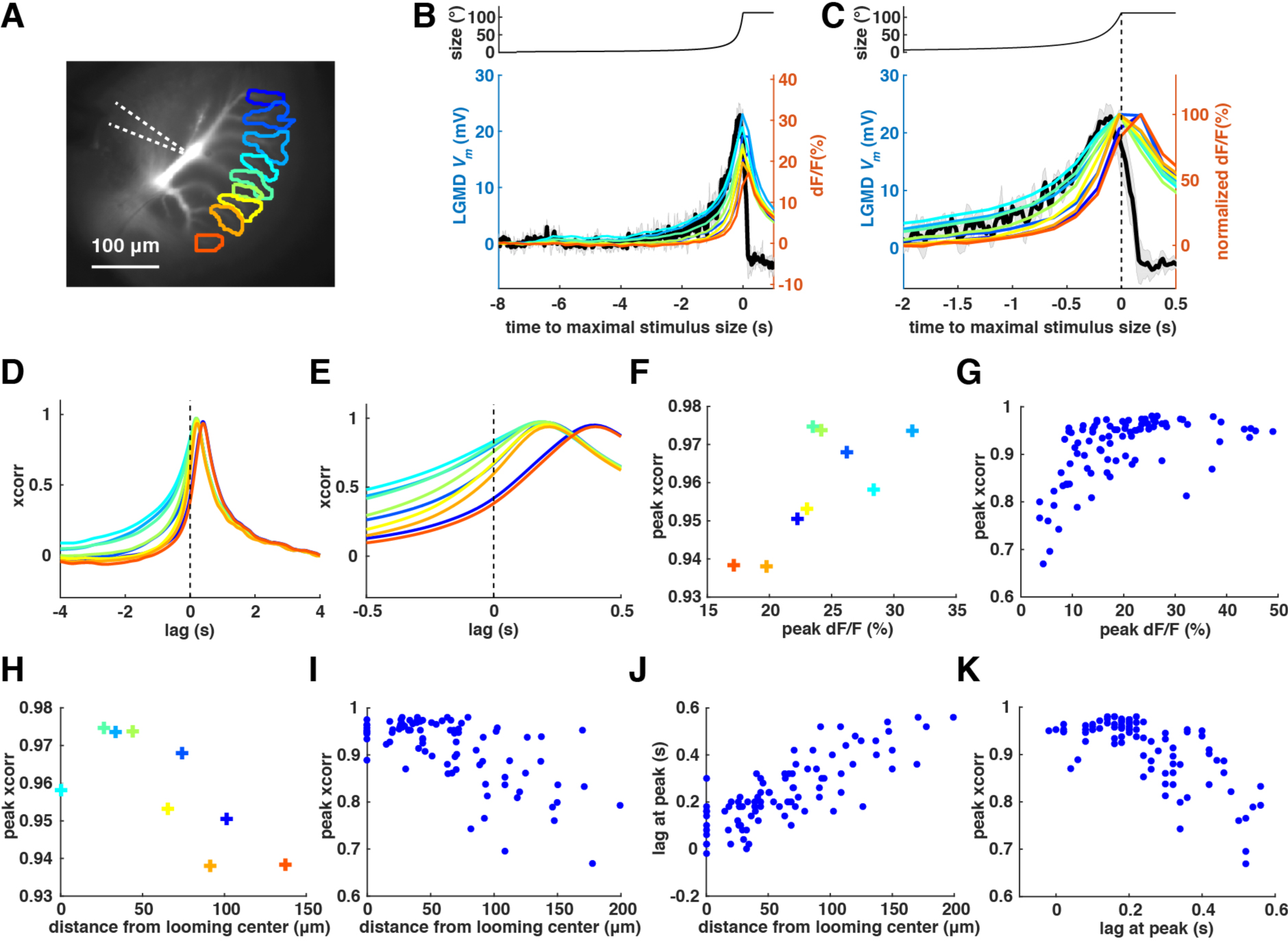
Comparison of the calcium responses and the subthreshold Vm. (A)Excitatory dendritic field of the LGMD with selected dendritic branches indicated by different colors. White dashed lines indicate the location of the intracellular sharp electrode. (B)***Top***: time course of the angular size of the looming stimulus with *l/|v|* = 120 ms. ***Bottom***: the dendritic membrane potential (Vm) of the LGMD is shown as the black solid trace. Gray shading, standard deviation of 4 trials. Thin solid colored traces are the time courses of the average dF/F within the selected dendritic branches in (A). (C)Zoomed-in view of (B). The dF/F traces were normalized to the peaks of each individual trace. (D)Cross-correlation (xcorr) of the rising phases of *V*_*m*_ and dF/F. Colors match those in (B). Black dashed line marks the zero-time lag. (E)Zoomed-in view of (D). (F)The peak of each cross-correlation trace in (D) plotted as a function of the peak dF/F for each corresponding dendritic branch in (B). (G)Same as (F) but across animals. (n=7 animals, Vm recorded at 11 locations). (H)The peak of each cross-correlation trace in (D) plotted as a function of the distance between the corresponding dendritic branch and the looming center. (I)Same as (H) but across animals. (n=7 animals, Vm recorded at 11 locations). (J)The time lag at the peak of each cross-correlation trace plotted across animals as a function of the distance between the corresponding dendritic branch and the looming center. (n=7 animals, Vm recorded at 11 locations). (K)The peak cross-correlation plotted as a function of the time lag at the peak across animals. (n=7 animals, Vm recorded at 11 locations).

We further analyzed the calcium responses on a finer spatial scale by considering each imaged dendritic branch separately (Fig. 1D, top). We first defined a threshold dF/F activation as 5 times the baseline noise of dF/F (dashed line, Fig. 1D, bottom; Methods). The baseline noise was defined as the maximum of the dF/F values over the 2 s preceding the onset of the visual stimulus. We call the time at which dF/F crosses this threshold relative to the time at which the stimulus reaches its maximal size the rise time. We compared the rise times of all the branches and found that the branch that had the earliest rise time mapped to the center square on the display (Fig. 1D and E, top; compare with Fig. 1A, bottom). In the following, we will call this branch the (dendritic) looming center. As predicted from retinotopy, the rise time increased in an orderly manner with the distance (in the imaging plane) of each branch from the looming center, a result consistent across animals (Fig. 1E, bottom). We also found that dF/F integrated over time up to its peak, an indirect measure of cumulative calcium influx, decreased as a branch lied farther from the looming center (Fig. 1F, top). This result was again consistent across animals (Fig. 1F, bottom). Thus, looming stimuli generate an orderly activation of excitatory dendritic branches that lasts over the entire course of the stimulus and decreases in strength.

### Calcium responses at looming center branches closely track subthreshold membrane potential

Calcium fluorescence changes are usually much slower than membrane potential (*V*_*m*_) changes due to the buffering properties of calcium dyes and their binding kinetics. In preliminary experiments, we compared the calcium responses to looming stimuli obtained using either OGB-1 or OGB-5N and found OGB-5N to have faster kinetics (Supp. Fig. 2A, B). Next, we checked how closely OGB-5N calcium responses track the subthreshold membrane potential by recording them simultaneously during the presentation of looming stimuli with *l/|v|* = 120 ms. The calcium signals from individual dendritic branches were analyzed separately (Fig. 2A) and are plotted together with *V*_*m*_ (median filtered to eliminate spikes) in Fig. 2B and on an expanded time base in Fig. 2C (black traces). The calcium responses on the looming center branches best matched the subthreshold *V*_*m*_. This was also true at faster approach speeds, when *l/|v|* = 30 and 60 ms (Supp. Fig. 2C-F). The match was particularly close over the rising phase of *V*_*m*_ up to its peak, when the synaptic excitation generating the calcium influx dominates, but not over the decaying phase of *V*_*m*_, when synaptic inhibition dominates (Gabbiani et al. 2002; Gabbiani et al. 2005). To quantify the relation between *V*_*m*_ and calcium fluorescence, we next computed the cross-correlation between the rising phase of *V*_*m*_ and that of the dF/F and plotted it as a function of the time lag in Fig. 2D and on an expanded time base in Fig. 2E. Across branches, we found high peak cross-correlation values that increased as the peak dF/F increased and eventually saturated (Fig. 2F, single animal; 2G across animals). In parallel, the peak of the cross-correlation decreased with distance from the looming center, further confirming that *V*_*m*_ was most closely linked to the looming center response (Fig. 2H, single animal; 2I across animals). Next, to finely resolve the spread of peak activation times across dendritic branches (c.f. Supp. Fig. 1B), we plotted the time lag at peak cross-correlation as a function of the distance between the considered branch and the looming center and found an orderly increase (Fig. 2J). The minimum lag at the looming center amounted to 0.11 ± 0.09 s (n = 7 animals). As expected from these last two results, the peak of the cross-correlation decreased with increasing time lag at the peak (Fig. 2K). To summarize, the calcium signal at the looming center closely tracked the membrane potential time course, with a delay of 110 ms; other branches experience nearly identical activation patterns that are increasingly delayed with distance from the looming center.

### Muscarine increases and scopolamine decreases the activation of the LGMD excitatory dendrites in response to looming stimuli

To study the effects of putative muscarinic acetylcholine receptors (mAChRs) located at the presynaptic terminals of the LGMD on looming responses, we compared the activation pattern on the LGMD excitatory dendritic field before and after puffing the agonist muscarine. As illustrated in one example in Fig. 3A, we found that the activated dendritic area increased after puffing muscarine. Calcium influx was also increased close to the spike initiation zone (SIZ; Fig. 3A, arrowhead), where voltage-gated calcium channels are activated by LGMD spiking (Peron and Gabbiani 2009), a result consistent with increased firing of the LGMD. When we compared the activation strength on a selection of dendritic branches (Fig. 3B), we found that muscarine increased it substantially (Fig. 3C, vs. 3D). This result was consistent across animals (Fig. 3E). To quantify the increase in area observed in Fig. 3A across animals, we computed the two principal axes of the activated area relative to its center of mass and projected the fluorescence pattern intensity along each of them (Supp. Fig. 3A-E). This procedure allowed us to obtain the full width at half maximum (FWHM) of the activated area along the principal axes (Supp. Fig. 3F) before and after drug application. As illustrated in Fig. 3F across five animals, there was a significant broadening (p=0.0002, paired t-test) of these curves following muscarine application. Next, we pooled the data across five animals and found that the peak dF/F decreased with increasing distance from the looming center both before and after puffing muscarine (Fig. 3G). Additionally, the magnitude of the slope of the peak dF/F as a function of the distance from the looming center significantly decreased after puffing muscarine.

We then compared the activation pattern on the LGMD excitatory dendritic field in response to looming stimuli before and after puffing the muscarinic antagonist scopolamine (Fig. 3H-N). As illustrated in one example in Fig. 3H, we found that the activation strength strongly decreased. Calcium influx close to the SIZ was also decreased (Fig. 3H, arrowhead), consistent with decreased firing of the LGMD. We next compared the dF/F on selected dendritic branches (Fig. 3I), and found a uniform decrease for all branches (Fig. 3J, vs. 3K). This result was consistent across animals (Fig. 3L), although the change in the activation area observed in the example of Fig. 3H was not significant after pooling across animals (Fig. 3M and legend). The peak dF/F also decreased with increasing distance from the looming center after puffing scopolamine (Fig. 3N). However, we did not observe a significant difference in the slope of the peak dF/F as a function of the distance from the looming center before and after puffing scopolamine (Fig. 3N). Thus, the effects of scopolamine are nearly opposite to those observed when applying muscarine.

Finally, we tested the effect of scopolamine on LGMD spiking by recording extracellularly its activity to small (1 × 1° and 2 × 2°) transient stimuli flashed at various positions across the visual field (Supp. Fig. 3G-I). On average, scopolamine decreased spiking by approximately 50%, further confirming the excitatory role of mAChRs and suggesting that they are tonically active since these small flashes likely activated one or two ommatidia at most. Taken together, these results strongly suggest that the mAChRs between adjacent presynaptic LGMD terminals are excitatory and that they significantly contribute to increasing synaptic calcium influx in response to looming stimuli.

### Scopolamine and muscarine do not affect dynamic lateral inhibition presynaptic to the LGMD

The LGMD’s response to a small translating object is inhibited by wide-field drifting stripes flanking it, provided they are sufficiently close (O’Shea M and Rowell 1975, Rowell et al. 1977). This dynamic lateral inhibition protects the LGMD collision-detection circuit from habituation caused by background motion (e.g., during locomotion) and plays a role in suppressing excitation early during a looming stimulus (Gabbiani et al. 2002; Rind et al. 2016). The location of this type of dynamic lateral inhibition along the pathway leading from photoreceptors to the LGMD remains unknown. Given the excitatory signature of mAChRs presynaptic to the LGMD, it appeared unlikely that they could implement dynamic lateral inhibition as well. However, since dynamic lateral inhibition has been predicted to be mediated by mAChRs in presynaptic terminals to the LGMD (Rind and Simmons, 1998, Rind et al. 2016), we designed visual stimuli to explicitly probe this possibility. The baseline condition consisted of a small translating black square that appeared at one edge of the screen 2 s after the start of the trial (Fig. 4A; Supp. Movie 2). The square remained stationary for 4 s and then moved at a speed of 14 °/s for 8 s until it completely moved outside of the display. This allowed us to separate the LGMD’s responses to the square’s appearance and its motion. In the test conditions, the same stimulus moved flanked by two gratings drifting in the same direction at a temporal frequency of 6 Hz (Fig. 4B). The distance between the two gratings was varied from 7.5 to 70° (0 to 31.25° between the edges of the moving square and the gratings) yielding maximal and minimal activation of dynamic lateral inhibition, respectively.

**Fig. 4.**
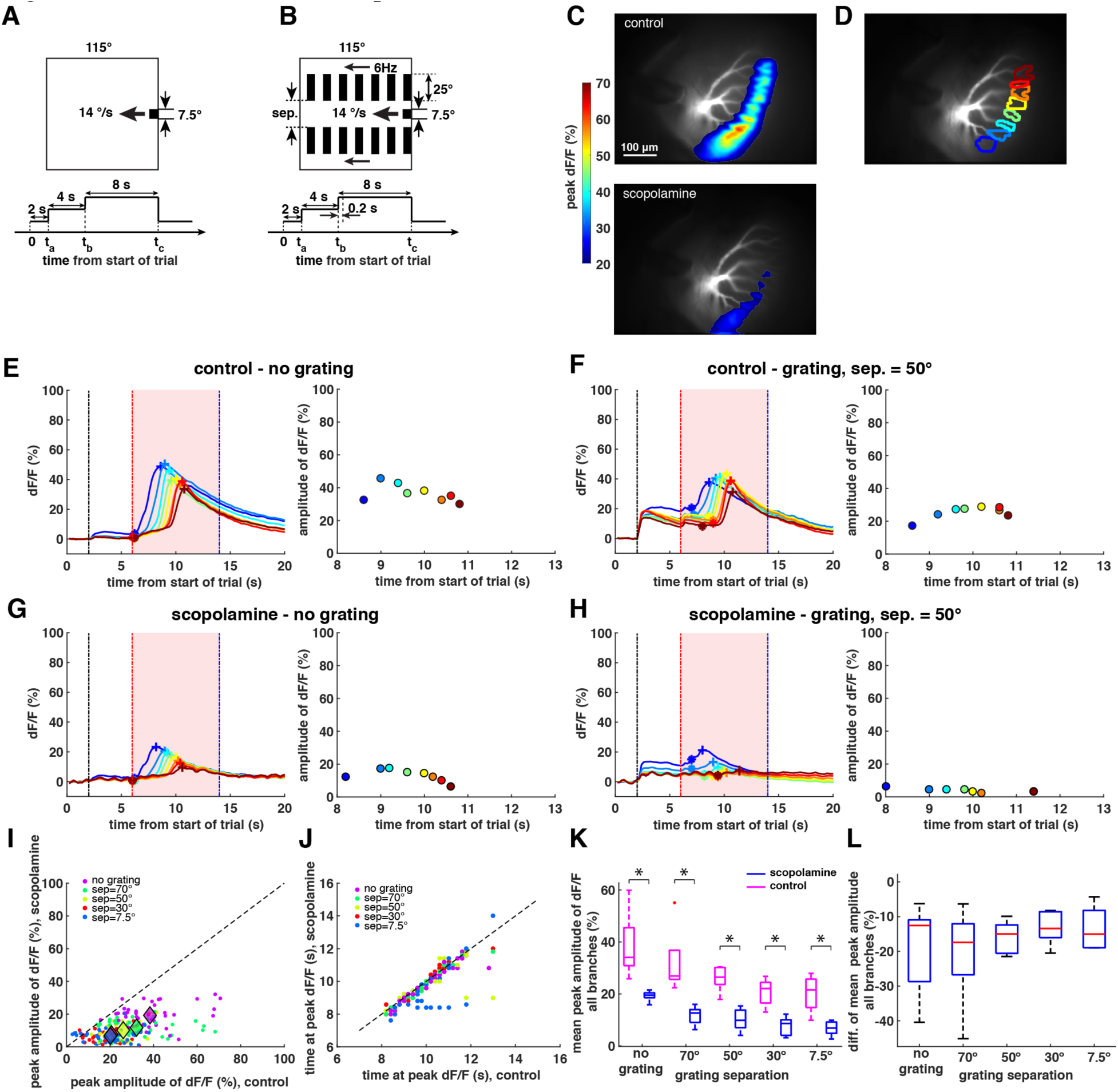
Test of the influence of scopolamine on lateral inhibition. (A) ***Top***: schematics of a small translating visual stimulus that moves at a speed of 14°/s. ***Bottom***: time course of the visual stimulus. Time 0 to t_a_: bright display; t_a_: small black stimulus flashed on at the right edge of the display; t_a_ to t_b_: stimulus stays still; t_b_ to t_c_: small black stimulus translates from the right to the left display edge until it completely moves outside of the display. (B) ***Top***: schematics of a small translating visual stimulus at a speed of 14°/s with two flanking gratings drifting laterally (temporal frequency: 6 luminance transitions per second, 90 º/s). The width of each grating is 25°. Abbreviation, sep.: separation. ***Bottom***: time course of the visual stimulus. Time 0 to t_a_: bright display; t_a_: small black stimulus flashed on at the right edge of the display, simultaneously the two gratings are flashed on; t_a_ to t_b_: stimulus stays still; t_b_ to t_c_: gratings start drifting; t_b_+0.2 s to t_c_: small black stimulus translates from the right to the left edge of the display until it completely moves outside of the display; t_c_: gratings stop drifting and stay still. (C) ***Top***: dendritic activation in response to a small translating visual stimulus. ***Bottom***: dendritic activation in response to the same small translating visual stimulus after puffing scopolamine. The small translating visual stimulus is the same as depicted in Fig. 4A. (D) Excitatory dendritic field of the LGMD with selected dendritic branches indicated by different colors. (E) ***Left***: time courses of the average dF/F at each of the selected dendritic branches in (B), with matched colors, in response to the same small translating visual stimulus as in Fig. 4A. Each trace is an average of dF/F over 3 trials. Black vertical dashed line: time when the small visual stimulus appears. Red vertical dashed line: time when the small visual stimulus starts translating. Blue vertical dashed line: time when the small visual stimulus moves completely outside of the display. The asterisk and plus markers on each curve indicate the baseline and the peak dF/F of the calcium response to the translating stimulus, respectively. ***Right***: the peak amplitude of dF/F is computed by subtracting the baseline (indicated by the asterisk marker on ***left***) from the peak dF/F (indicated by the plus marker on ***left***) for each curve on ***left*** with matched color and plotted as a function of the time at peak dF/F. (F) ***Left***: time courses of the average dF/F (over 3 trials) at each of the selected dendritic branches in (B), with matched colors, in response to a small translating visual stimulus with lateral drifting gratings (with a separation between gratings of 50°) as in Fig. 4B. Black vertical dashed line: time when the small visual stimulus and lateral gratings appear. Red vertical dashed line indicates the time when the lateral gratings starts drifting and 0.2 s before small visual stimulus starts translating. Blue vertical dashed line: time when the small visual stimulus moves completely outside of the display and the lateral gratings stop drifting. The asterisk and plus markers on each curve indicate the baseline and the peak dF/F of the calcium response to the translating stimulus, respectively. ***Right***: same plotting conventions as in (C) ***right***. (G) Same visual stimuli and same plotting conventions as in (C) after puffing scopolamine. (H) Same visual stimuli and same plotting conventions as in (D) after puffing scopolamine. (I) The amplitude of dF/F on every selected dendritic branch after puffing scopolamine versus before puffing (5 animals, 41 dendritic branches total, average of 3 trials per condition). Blue dots: small translating stimuli with no lateral drifting gratings. Green, brown, red, and purple dots: small translating stimuli with lateral drifting gratings with separations of 70°, 50°, 30° and 7.5°, respectively. Purple, green, light green, red and blue diamonds are the mean (center-of-mass) of the purple, green, light green, red and blue dots, respectively. (J) Time at peak dF/F on every selected dendritic branch after puffing scopolamine versus before puffing. Same plotting conventions as in (I). (K) Box plots of the mean peak amplitudes of dF/F for all branches in each animal in response to small translating stimuli with no lateral drifting gratings, and with gratings separated by 70°, 50°, 30° and 7.5°, respectively. Blue, after puffing scopolamine. Magenta, before puffing scopolamine. (n=5 animals, * p<0.05, Wilcoxon signed-rank test, uncorrected for multiple comparisons; p=0.025 for control and p=0.0001 for scopolamine, one-way ANOVA; p=0 for control vs. scopolamine, two-way ANOVA). (L) Box plots of the difference of mean peak amplitudes for all branches in each animal after puffing scopolamine and before puffing in response to small translating stimuli with no lateral drifting gratings, and with gratings separated by 70°, 50°, 30° and 7.5°, respectively. No significant statistical difference of the means was found between any two groups. (n=5 animals, p=0.66 one-way ANOVA).

We first tested the influence of scopolamine on dynamic lateral inhibition. Interestingly, in response to the small translating stimulus the dendritic area activated under control conditions was larger than that activated by looming stimuli (compare Fig. 4C, top with Figs. 3A, 3H). This is surprising as looming stimuli elicit much stronger firing in the LGMD than translating stimuli (Peron and Gabbiani, 2009). We will elaborate on this observation further below. After puffing scopolamine, dendritic activation decreased (Fig. 4C, bottom). On each selected dendritic branch (Fig. 4D), the amplitude of dF/F in response to a small translating stimulus was stronger than in response to the same stimulus flanked by drifting gratings at a separation of 50°, consistent with activation of lateral inhibition (Fig. 4E, 4F). After puffing scopolamine, the amplitude of the dF/F on each selected dendritic branch decreased (Fig. 4G, 4H). However, the lateral inhibition persisted. We employed flanking gratings with separations of 70, 50, 30 and 7.5° and computed the amplitude of dF/F on every selected dendritic branch after puffing scopolamine versus before puffing in five animals (Fig. 4I). The amplitudes of dF/F after puffing scopolamine were significantly smaller than before puffing across the animals (p<10^−9^ for all stimuli types, paired t-test). The time at peak dF/F before and after puffing scopolamine did not significantly change for two separations (30° and 50°) while in the remaining three cases the change was small (-1.1% change at 70°; −4.2% at 7.5° and −1.8% for no grating; Fig. 4J). The mean amplitudes of dF/F for all the branches in each animal decreased with closer flanking gratings, both before and after puffing scopolamine (Fig. 4K). However, the subtraction of the mean amplitudes of dF/F before puffing scopolamine from that after puffing showed no statistical difference in response to all five different visual stimuli (Fig. 4L and legend). This suggests that blocking mAChRs presynaptic to the LGMD with scopolamine did not influence dynamic lateral inhibition.

Next, we tested the influence of muscarine on dynamic lateral inhibition using the same protocol as above. In response to the small translational stimulus, activation on the LGMD excitatory dendrites increased after puffing muscarine (Supp. Fig. 4A). On each selected dendritic branch (Supp. Fig. 4B), we compared the amplitude of dF/F in response to the small translational stimulus without and with the drifting gratings at a separation of 50° (Supp. Fig. 4C and D, respectively). We found that the overall amplitude of dF/F was reduced in presence of the gratings, consistent with the activation of presynaptic lateral inhibition. After puffing muscarine, the amplitude of the dF/F on each selected dendritic branch increased (Supp. Fig. 4E and F). However, the responses with gratings were still reduced compared to the no grating condition. Thus, lateral inhibition appeared to be unaffected by activation of the mAChRs in presynaptic terminals to the LGMD. We repeated this experiment across five animals, using gratings with separations of 70, 50, 30 and 7.5° and plotted the amplitude of dF/F on every selected dendritic branch after puffing muscarine versus before puffing (Supp. Fig. 4G). The amplitudes of dF/F after puffing muscarine were slightly but not significantly bigger than before puffing across the animals. The time at peak dF/F before and after puffing muscarine did not change (Supp. Fig. 4H). The mean amplitudes of dF/F across all the branches in each animal decreased as the distance between the drifting gratings decreased both before and after puffing muscarine, the characteristic signature of lateral inhibition (Supp. Fig. 4I). However, subtraction of the mean amplitudes of dF/F before puffing muscarine from that after puffing showed no statistical difference between the five different visual stimulus conditions (Supp. Fig. 4J). Thus, muscarine like scopolamine did not affect dynamic lateral inhibition.

Taken together, lateral interactions between transmedullary afferents through mAChRs at presynaptic terminals onto the LGMD do not appear to generate dynamic lateral inhibition.

### Horizontal band-restricted looming stimuli activate lateral branches better than looming stimuli

To further study the dendritic activation pattern elicited by looming stimuli, we designed looming stimuli restricted to horizontal (along the dorsal-ventral eye axis due to the 90° head rotation performed during the dissection; Methods) or vertical (along the posterior-anterior eye axis) bands with a width of 8° (Fig. 5A). As expected from the retinotopic mapping, the activation width of the vertical band-restricted looming response (Fig. 5A, right) along the first principal axis (magenta line) was narrower than that of the standard looming response (Fig. 5A, left). Surprisingly, we found that the activation width of the horizontal band-restricted looming response (Fig. 5A, middle) was wider than that of the standard looming response (Fig. 5A, left). On each selected dendritic branch (Fig. 5B), we compared the calcium responses to the standard looming stimulus (Fig. 5C, left), the looming stimulus restricted to the horizontal band (Fig. 5C, middle) and to the vertical band (Fig. 5C, right), respectively. We found that the farther a branch was from the looming center the smaller its calcium responses were both to standard looming (Fig. 5C, left) and to vertical band-restricted looming (Fig. 5C, right). The latter responses decreased even more strongly with distance from the looming center. However, in response to horizontal band-restricted looming, calcium responses did not decrease with distance from the looming center and even increased slightly (Fig. 5C, middle). To show this more directly, we plotted the peak dF/F on each selected dendritic branch, normalized to the peak dF/F on the branch that mapped to the looming center, as a function of distance to the looming center across animals (n=5 animals; Fig. 5D). We found that with increased distance from the looming center the normalized peak dF/F decreased fastest in response to the vertical band-restricted looming stimulus, while the normalized peak dF/F increased a bit and then decreased slowest in response to the horizontal band-restricted looming stimulus (Fig. 5D). By comparing the activated width along the first principal and second principal axis in response to standard looming, to horizontal, and to vertical band-restricted looming across animals (Fig. 5E), we found that the activated widths along the second principal axis for the three cases were similar (blue traces; p=0.15, one-way ANOVA). However, along the first principal axis the activated width in response to horizontal band-restricted looming was the biggest, and the activated width in response to vertical band-restricted looming was the smallest (red traces). Interestingly, along the first principal axis, the activation strength at the center branch is not the strongest in response to horizontal band-restricted looming as there is a clear groove at the center position (Fig. 5E, middle). We computed the ratio of the full-width-at-half-maximum (FWHM) of the activation along the first principal axis in response to horizontal or vertical band-restricted looming to that in response to standard looming (Fig. 5F). The horizontal-based ratio was larger than 1 and the vertical-based ratio smaller than 1 for all the animals (Fig. 5F, n=5, p=0.0088, paired t-test). When we carried out a similar analysis for small translating squares motivated by the wide fluorescence pattern observed in Fig. 5C, we also found increased activation for small squares relative to looming stimuli (Supp. Fig. 5).

**Fig. 5.**
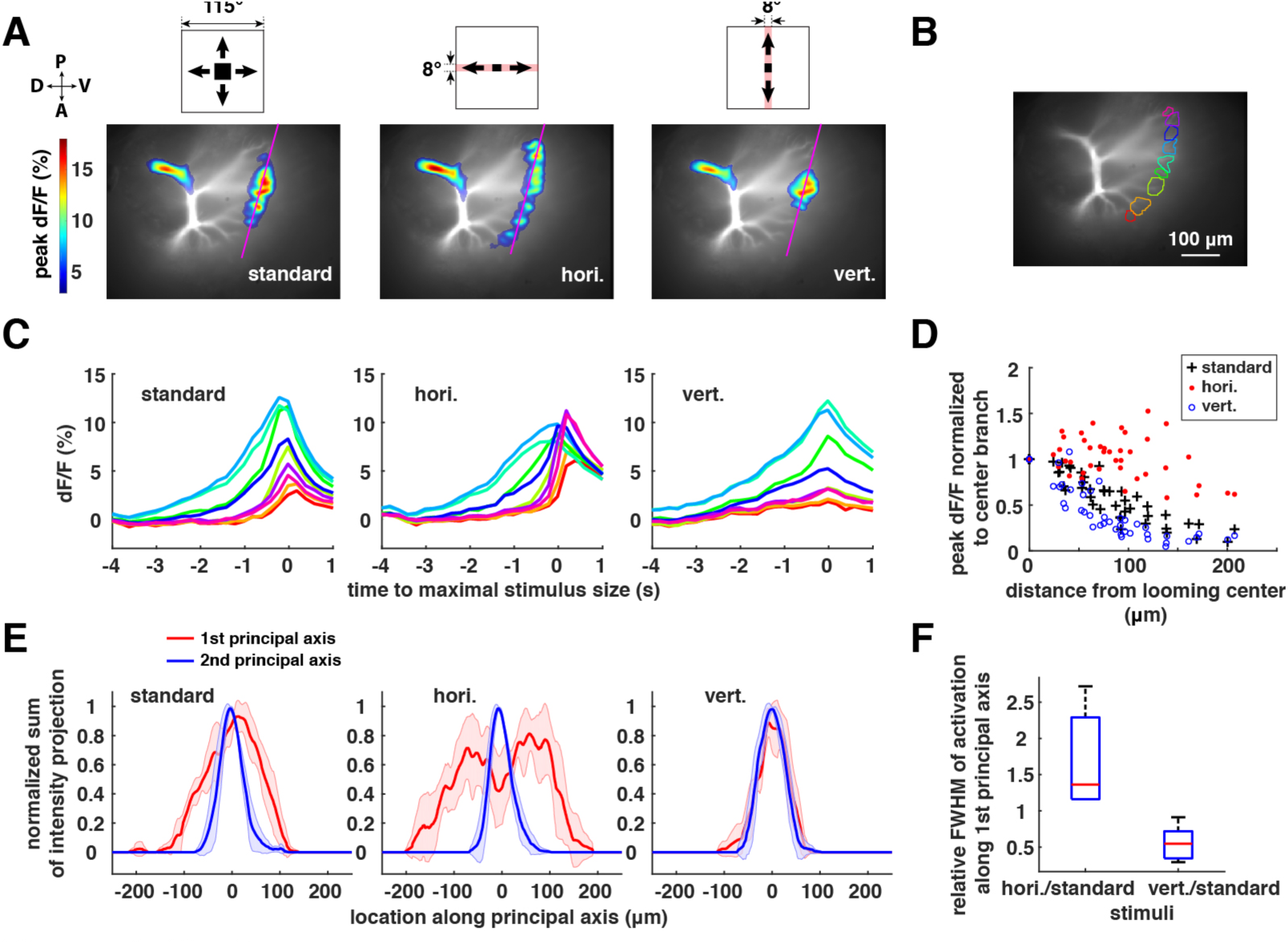
Calcium responses to looming stimuli restricted to horizontal and vertical bands. (A) ***Top:*** from left to right, schematics of a standard looming stimulus, looming stimulus restricted in a horizontal band (along the dorsal-ventral eye axis) and looming restricted in a vertical band (along the posterior-anterior eye axis), respectively. ***Bottom***: from left to right, activation on the LGMD excitatory dendrite in response to the looming stimulus, looming restricted in a horizontal band and looming restricted in a vertical band, respectively. (B) Excitatory dendrite of the LGMD with selected dendritic branches indicated by different colors. (C) From left to right, time courses of the average dF/F at each of the selected dendritic branches in (B), with matched colors, in response to the looming stimulus, looming restricted in a horizontal band and looming restricted in a vertical band, respectively. (D) Peak dF/F of each branch normalized to the peak dF/F of the center branch and plotted as a function of the distance from the branch that mapped to the looming center. Black crosses, responses to standard looming; Red dots, responses to looming restricted in a horizontal band; Blue circles, responses to looming restricted in a vertical band. (E) Sum of peak dF/F projected onto the 1^st^ and 2^nd^ principal axes and plotted as a function of the projection location along that principal axis after normalizing to the peak of the curve. Red, along first principal axis; Blue, along second principal axis. Red and blue shadings, standard deviation of 5 animals. ***Left, middle and right***, response to a standard looming stimulus, looming restricted in a horizontal band and looming restricted in a vertical band, respectively. (F) Ratio of the full-width at half-maximum (FWHM) of the curve in (E) along the first principal axis in response to the horizontal/vertical band-restricted looming stimulus to the FWHM of the curve in (E) along the first principal axis in response to the standard looming stimulus. (n=5 animals).

### Presynaptic mAChRs partially counteract global inhibition upstream of the LGMD

To better understand why the activation of lateral branches is weaker in response to standard looming stimuli than to band-restricted horizontal looming stimuli and small translating squares, we designed flicker-off stimuli with different angular sizes (Fig. 6A). The activated dendritic area initially increased as the size of the stimulus increased (Fig. 6B). However, for sizes exceeding 40° the magnitude of the peak dF/F decreased dramatically as angular size further increased up to 100° (Fig. 6B). This suggests the existence of size-dependent inhibition upstream of the LGMD excitatory dendrites that was strongly activated by stimuli larger than 40º. Next, we tested the influence of scopolamine and muscarine on this novel, global type of inhibition. The peak dF/F of the center branch (Fig. 6C) and the mean peak dF/F of all selected branches (Fig. 6D) decreased as the size of the stimulus exceeded 40°. After puffing scopolamine, the peak dF/F of the center branch (Fig. 6C) and the mean peak dF/F of all selected branches (Fig. 6D) was reduced for all stimuli, irrespective of size. After puffing muscarine, the peak dF/F of the center branch (Fig. 6E) and the mean peak dF/F of all the selected branches (Fig. 6F) increased as the size of the stimulus increased from 10° to 40°, but exhibited no clear decrease as the size of the stimulus further increased to 100°. These results suggest that mAChRs partially counteract the effect of global inhibition upstream of the LGMD that is triggered by large visual stimuli.

**Fig. 6.**
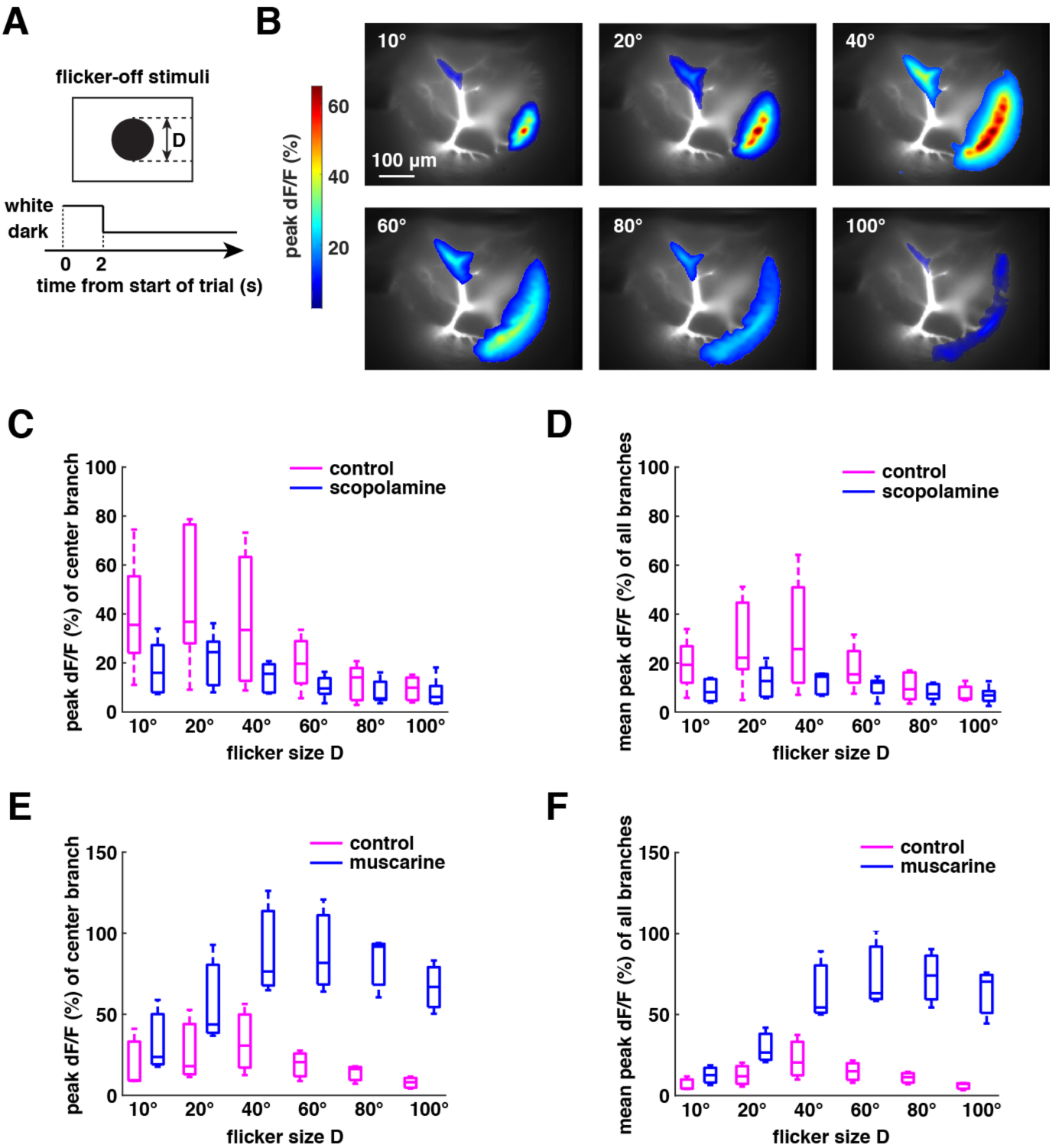
Influence of scopolamine and muscarine on calcium responses to flicker-off stimuli with different sizes. (A) ***Top***: schematics of flicker-off stimuli with a diameter of D. ***Bottom***: time course of the visual stimulus. Time 0 to 2 s: bright display; time after 2 s: black stimulus flashed on at the center of the display. (B) activation of the LGMD in response to flicker-off stimuli with sizes of 10°, 20°, 40°, 60°, 80° and 100°, respectively. (C) Boxplot of the peak dF/F for the branch that mapped to the stimulus center in response to flicker-off stimuli with sizes of 10°, 20°, 40°, 60°, 80° and 100°, respectively. Magenta and blue are without and with scopolamine, respectively. (n=5 animals; p=0.0007 for control vs. scopolamine, two-way ANOVA; p=0.028 for control, p=0.069 for scopolamine, one-way ANOVA). (D) Boxplot of the mean peak dF/F over all branches in response to flicker-off stimuli with sizes of 10°, 20°, 40°, 60°, 80° and 100°, respectively. Magenta and blue are without and with scopolamine. (n=5 animals; p=0.0009 for control vs. scopolamine, two-way ANOVA; p=0.07 for control, p=0.35 for scopolamine, one-way ANOVA). (E) Boxplot of the peak dF/F for the branch that mapped to the stimulus center in response to flicker-off stimuli with sizes of 10°, 20°, 40°, 60°, 80° and 100°, respectively. Magenta and blue are without and with muscarine. (n=3 animals; p=0 for control vs. muscarine, two-way ANOVA; p=0.45 for control, p=0.13 for muscarine, one-way ANOVA). (F) Boxplot of the mean peak dF/F over all branches in response to flicker-off stimuli with sizes of 10°, 20°, 40°, 60°, 80° and 100°, respectively. Magenta and blue are without and with muscarine, respectively. (n=3 animals; p=0 for control vs. muscarine, two-way ANOVA; p=0.15 for control, p=0.00004 for muscarine, one-way ANOVA).

### Blocking muscarinic lateral excitation decreases the LGMD’s spatial coherence preference

Recent research has demonstrated that the LGMD is selective for the spatial coherence of approaching objects and that this selectivity is partially explained by the interaction of hyperpolarization activated, cyclic nucleotide gated (HCN) channels with a slowly inactivating K^+^ current in the dendrites of the LGMD (Dewell and Gabbiani, 2017). Since lateral excitation mediated by mAChRs presynaptic to the LGMD is necessarily local based on anatomical and electrophysiological measurements (Rind et al. 2016; Ying and Gabbiani, 2016), we reasoned that it might also help increase responses to coherently expanding objects as opposed to incoherent ones. To test this hypothesis, we designed incoherent stimuli by first dividing the screen in a regular array of 2° wide ‘coarse pixels’ roughly spanning the receptive field of a single ommatidium. We replaced the dark edge motion associated with a looming stimulus expanding across each coarse pixel by an equivalent uniform decrease in luminance. The resulting stimulus, which we call a ‘coarse loom’ (Fig. 7A, middle), elicits responses in the LGMD that are indistinguishable from those of standard looming stimuli (Jones and Gabbiani 2010; Dewell and Gabbiani, 2017). Stimuli with decreasing degrees of coherence can then be generated from coarse looms by randomly displacing individual coarse pixels increasingly far from their original position (Fig. 7A, bottom).

**Fig. 7.**
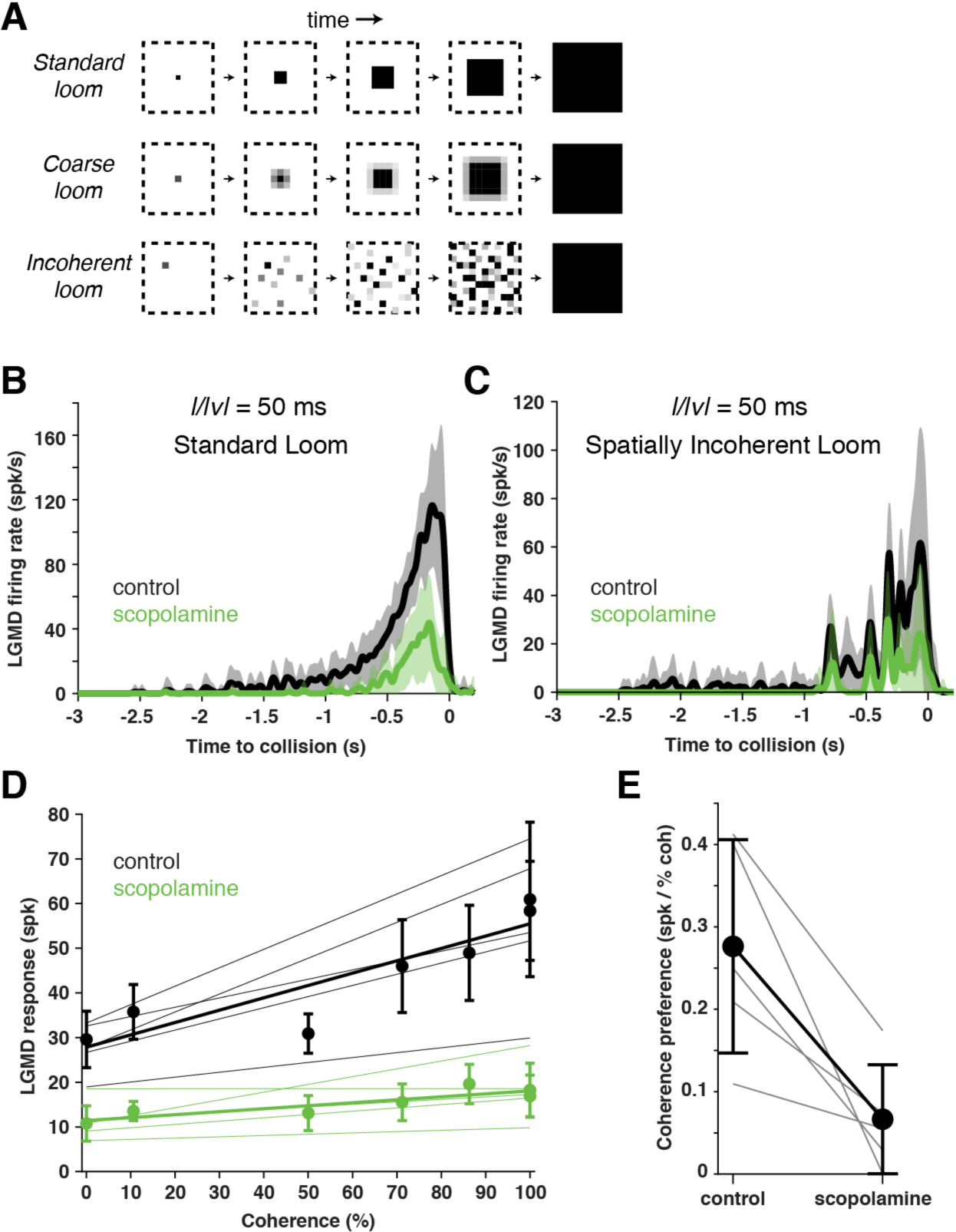
Scopolamine reduced looming responses and coherence preference. (A) Illustration of coherent and incoherent stimuli. For coarse looms (middle row) grayscale levels are set so that luminance in each coarse pixel is equal to that of standard looms in every frame. For reduced coherence looms (bottom), the spatial locations of the coarse pixels were altered (see Methods). (B) LGMD’s response to looming stimuli (*l/|v|* = 50 ms) decreased after addition of scopolamine. Lines and shaded region are mean and ± 1 sd (n = 5). (C) LGMD’s response to a spatially randomized version of a looming stimulus also decreased after addition of scopolamine. Lines and shaded region are mean and ± 1 sd (n = 5). (D) LGMD responses increase with stimulus coherence (Pearson r=0.90, p=0.006), but this coherence preference decreased after scopolamine (p = 0.008, ANCOVA test of slopes; n=5). Thin lines are data from individual animals. Thick lines and points are population average. Error bars are ± 1 sd. (E) Scopolamine caused a decrease in coherence preference (calculated from the slopes of the lines in C) p = 0.022, paired t-test. On average, there was a decrease from 0.28 spikes per % coherence to 0.07 spikes per % coherence.

To test whether mAChR-mediated lateral excitation increases coherence preference, we presented such stimuli while recording the firing rate of the LGMD before and after blocking it with scopolamine. Across stimulus conditions, the timing of LGMD firing remained similar after adding scopolamine, but with much lower firing rates (Fig. 7B, C). Responses to standard looming stimuli decreased from 58±16 spikes to 17±7 spikes after addition of scopolamine (mean±sd; p = 0.004). Responses to incoherent stimuli were also reduced after scopolamine addition (Fig. 7C), with 30±8 spikes in control and 11±6 spikes after puffing (p=0.017). For every animal tested there was a decrease in LGMD firing (p≤0.004) and the coherence dependent increase in firing was reduced (Fig. 7D). The coherence preference decreased for each animal (n=5) and on average there was a decrease from 0.28 spikes per percent coherence to 0.07 spikes per percent coherence (Fig. 7E). As scopolamine reduced responses to stimuli of all coherence levels, we calculated the coherence specific reduction by subtracting from the reduction in coherent looming response the reduction in incoherent response, and expressing it as a percentage of the control looming response. We found a 37.1 ± 17.8 % specific reduction for responses to coherent looming stimuli (p=0.01, paired t-test). These results indicate that lateral excitation indeed plays a role in shaping the LGMD’s preference for coherently expanding stimuli.

## Discussion

Little is known about how networks converging onto collision-detecting neuron are wired to shape their selectivity. Using calcium imaging, we investigated experimentally (for the first time to the best of our knowledge) how network connectivity among presynaptic excitatory inputs to a collision-detecting neuron shapes its responses. We provide evidence for two new types of presynaptic connectivity patterns: lateral excitation and global (normalizing) inhibition. We document one function of lateral excitation, which is to promote responses to coherently expanding stimuli and thus help tune the neuron to its preferred features. In this context, the presence of global inhibition fulfills the natural role of keeping excitation in check and within the dynamic range of the neuron. Together with the classical observations evidencing the role played by dynamic lateral inhibition in suppressing background motion noise (O’Shea and Rowell, 1975), these experiments show that, at least in the case of the LGMD, the presynaptic interactions among excitatory inputs to collision-detecting neurons are governed by a complex, but interpretable, set of rules.

Further, the results presented here also demonstrate that calcium imaging was effective at characterizing the spatiotemporal dynamics of excitatory synaptic activation on the LGMD dendrites in response to looming stimuli, and fast enough to match the subthreshold membrane potential dynamics recorded intracellularly in the excitatory dendritic field. Local application of both muscarine and scopolamine on the LGMD’s excitatory dendritic field yielded consistent results, establishing that local interactions among presynaptic LGMD afferents are excitatory. Notably, these excitatory interactions could not be evidenced in earlier experiments, where only two adjacent ommatidia were successively stimulated at intervals ranging from 17 ms to 1 s (Jones and Gabbiani, 2010; Ying and Gabbiani, 2016). This suggests that at least two adjacent ommatidia (and probably more) need to be stimulated to affect a third one.

Mammals possess five types of mAChRs (m1-m5), of which m1, m3 and m5 are coupled to G protein members of the G_q/11_ family, leading to activation of phospholipase C-beta and an intracellular IP3/Ca^2+^ cascade, which mostly results in excitatory effects. Two other mAChRs types (m2 and m4) are coupled to G protein members of the G_i/0_ family, which leads to presynaptic or postsynaptic inhibition (Brown 2010). In *Drosophila* and other arthropods three types of mAChRs have been found: types A, B and C (Xia et al. 2016). Type A structurally resembles most the vertebrate mAChRs, is activated by acetylcholine and muscarine, and is blocked by the classical mAChR antagonists, atropine and scopolamine. Type B is also activated by acetylcholine, but is not sensitive to muscarine, and is not blocked by the classical antagonists (Collin et al. 2013). The type A mAChR is coupled to G_q/11_ type proteins, while type B is coupled to G_i/0_ (Ren et al. 2015). The recently discovered type C resembles pharmacologically the type A mAChR (Xia et al. 2016). Our pharmacological results suggest that the mAChRs presynaptic to the LGMD are either of type A or type C. Their excitatory effects might result from inhibition of presynaptic M channels or other potassium channels (Delmas and Brown 2005; Shen et al. 2005; Brown 2010). In *Drosophila*’s visual system, small-field directional motion-selective T4 and T5 cells, transmedullary neurons that synapse onto lobula plate tangential cells, both express transcripts of nicotinic and muscarinic acetylcholine receptors (Shinomiya et al. 2014). Although the role of mAChRs in this system remains to be determined, our results suggest that the effects of their activation are likely to last at least 100 ms, a time scale that is long relative to that thought to be relevant for directional motion detection (Behnia et al., 2014). In vertebrates, mAChRs have been mainly associated with modulation of attention during behavior (e.g., Niell and Stryker, 2010). Our results suggest they may also contribute to visual detection tasks requiring local modulation of activity on a slow time scale, such as the role in object segmentation evidenced in this work.

The experiments carried out over the course of this work yielded several unexpected results. First, calcium activation consistently outlasted the time interval over which a looming stimulus swept across the retinotopic region associated with a given dendritic branch. This was particularly striking for the dendritic looming center, where relative fluorescence changes outlasted the stimulus presence by as much as 1 s (Fig. 1B, C). Two observations help explain this feature of the dendritic calcium response. First, the continued rise of calcium fluorescence immediately after the stimulus has left the red shaded region in Fig. 1B, C is caused in part by the overlapped dendritic activation for ommatidia up to 8° away (red double arrow in Fig. 1C; Ying and Gabbiani, 2016). Second, although the edges of the stimulus leave the retinotopic region associated with the dendritic looming center, its expanding vertical edges continue to stimulate the same dendritic branch and thus cause additional calcium entry there (Fig. 5A, C). Although muscarine and scopolamine affected the strength of calcium signals (Fig. 3), they did not significantly affect their time course, ruling out excitatory lateral interactions presynaptic to the LGMD as a major determinant of increasing responses beyond the region of dendritic overlap discussed above.

A second unexpected result was the excitatory nature of the mAChR-mediated lateral interactions presynaptic to the LGMD. Up to this point, the only demonstrated lateral interactions presynaptic to the LGMD along its excitatory projection were those of dynamic lateral inhibition (O’Shea and Rowell, 1975). It was thus plausible that they would be implemented by the lateral connections evidenced in electron microscopy reconstructions at the level of the LGMD (Rind and Simmons, 1998). Our results (Fig. 4, Supp. Fig. 4) indicate that this is not the case and that dynamic lateral inhibition therefore has to be located upstream of the LGMD, possibly even in the lamina or the medulla based on the broad pattern of GABA-ergic immunoreactivity observed in the locust optic lobe (Rosner et al., 2017).

A third unexpected result stems from the experiments summarized in Figs. 5, 6 and Supp. Fig. 5. They bring to evidence the presence of size-dependent, global inhibition onto presynaptic excitatory afferents located upstream of lateral excitation. Whether this global inhibition and dynamic lateral inhibition are generated from the same inputs is still unknown. Global inhibition explains why the dendritic activation width for a standard looming stimulus is smaller than that observed for a band-restricted looming stimulus or for a small translating stimulus. The broader function of global inhibition is most likely to normalize the strength of the excitatory inputs impinging onto the LGMD. As a looming stimulus expands, the number of activated ommatidia grows nearly exponentially (e.g., Fig. 1B, top) and the increase in edge velocity increases the strength of individual synaptic inputs as well (Jones and Gabbiani 2010). Without normalization, excitation would be expected to grow more than one thousandfold over the course of a looming stimulus. Thus, global inhibition would serve to normalize these inputs into the dynamic range of the LGMD. The results reported in Fig 1F are consistent with this hypothesis. A similar role for normalizing global inhibition has been documented in several other sensory systems both in vertebrates and invertebrates (review: Carandini and Heeger, 2012). After puffing muscarine onto the LGMD excitatory dendritic field, its responses to large flicker stimuli were enhanced and its dendritic activation width to looming stimuli was increased. This indicates that the activation of mAChRs tends to counteract global inhibition.

As lateral excitation is local, requires activation of several neighboring ommatidia, and occurs on a relatively slow time scale, this suggested to us that it might be involved in discriminating between coherently and incoherently expanding looming stimuli, a form of object segmentation. We confirmed this hypothesis experimentally by showing that coherence sensitivity was diminished in the output firing rate of the LGMD following block of lateral excitation (Fig. 7). Local lateral excitation is thus a mechanism that helps tune the LGMD to looming stimuli and ignore spatially incoherent ones, in addition to dendritic conductances located within the LGMD’s dendrites (Dewell and Gabbiani 2017). Although both pre- and post-synaptic mechanisms influence the spatial selectivity of LGMD responses, the coherence tuning due to presynaptic mechanisms shown here plays a somewhat smaller role than that of the dendritic conductances reported in our earlier work (median coherence-specific reduction of 57% after blocking HCN channels, computed from Dewell and Gabbiani, 2017, vs. 34% median specific reduction after scopolamine addition reported here).

We have summarized in Supp. Fig. 6 the new connections evidenced by our data in the presynaptic circuitry to the LGMD and contrasted them with the model thought to be adequate up to this point. These results have implications for other collision detection circuits. Particularly interesting will be to determine how they perform visual object segmentation and whether the biophysical mechanisms involved resemble those implemented in the locust visual system.

## Methods

### Preparation

Experiments were carried out on mature locusts (mostly female), 3-4 weeks past the final molt, taken from a crowded colony maintained at Baylor College of Medicine. Animals were mounted dorsal side up on a custom holder with their heads rotated 90° to make the anterior side point downward. The head and neck were bathed in ice-cold locust saline, except for the right eye used for visual stimulation. The head capsule was opened dorso-frontally between the two eyes. The gut and the muscles in the head capsule were removed to reduce brain movement. The head was carefully detached from the body, leaving only the two nerve cords and four tracheas attached. The right optic lobe was desheathed with fine forceps. A metal holder was placed underneath the brain and the right optic lobe to elevate and stabilize them. The dissection of the animals typically lasted 1.5 h.

### Visual stimulation

Visual stimuli were generated with the Psychtoolbox and Matlab (The MathWorks, Natick, MA). A Digital Light Processing projector module (DLP LightCrafter 4500, Texas Instruments, Dallas, TX) and a diffuser screen (Non-adhesive stencil film, 0.1 mm thick, Darice, Strongsville, OH) were used to provide the visual stimuli. The projector was programmed to work in the pattern sequence mode with 2-bit bitmap images in green color which achieved an effective refresh rate of 243 Hz. The brightness of the display before presenting the looming stimulus was 38.7 cd/m^2^. The screen brightness at the level used for "off" stimuli was 0.06 cd/m^2^. The brightness of the screen was calibrated by a photometer and linearized by loading a normalized gamma table to the Psychtoolbox. The screen was placed 25 mm away from the locust eye and measured 79-by-79 mm. For the experiments presented in Fig. 7, visual stimuli were generated using Matlab and the PsychToolbox (PTB-3) on a PC running Windows XP. A conventional cathode ray tube (CRT) monitor refreshed at 200 frames per second was used for stimulus display (LG Electronics). The monitor was calibrated to ensure linear, 6-bit resolution control over luminance levels. A 100-120 s delay was used between stimuli and locusts were repeatedly brushed and exposed to light flashes and high frequency sounds to decrease habituation. ‘Coarse’ looming stimuli were generated as in our earlier work (Jones and Gabbiani 2010; Dewell and Gabbiani 2017). Briefly, the stimulation monitor was first pixelated with a spatial resolution approximating that of the locust eye (2° x 2°), referred to as ‘coarse’ pixels. Each coarse pixel’s luminance followed the same time course as that elicited by the edge of the simulated approaching object sweeping over its area. To alter the spatial coherence of these stimuli, a random two-dimensional Gaussian jitter with zero mean was added to each coarse pixel screen location.

The jittered positions were rounded to the nearest available coarse pixel location on the screen to prevent any coarse pixels from overlapping. The standard deviation of the Gaussian was altered between 0 and 80° to control the amount of shifting and thus the resulting spatial coherence of the randomized stimulus.

### Staining with calcium indicators

At the beginning of an experiment, the LGMD neuron was impaled with a sharp intracellular electrode (230-300 MΩ) containing 3 µl of 2 M potassium chloride and 1 µl of 5 mM Oregon Green BAPTA-1 (OGB-1) or Oregon Green BAPTA-5N (OGB-5N; hexapotassium salt; Thermo Fisher Scientific, Waltham, MA). Due to the stronger signal amplitude of OGB-1 compared to OGB-5N, OGB-1 staining was used in all the experiments which investigated the effects of puffing of muscarine or scopolamine on the LGMD’s excitatory dendrites. For other experiments, we used OGB-5N as its kinetics is faster than that of OGB-1. Iontophoresis of OGB-1 or OGB-5N was achieved with current pulses of −3nA, alternating between 1 s on and 1 s off, that lasted for 6 min. The pulses were delivered by an Axoclamp 2B amplifier (Molecular Devices, Sunnyvale, CA). The LGMD was identified through its uniquely characteristic spike pattern in 1:1 correspondence with that of its postsynaptic target, the descending contralateral movement detector (DCMD; O’Shea and Williams, 1974), recorded extracellularly from the nerve cord with hook electrodes.

### Imaging, electrophysiology, pharmacology, and data acquisition

We employed wide-field calcium imaging during visual stimulation experiments. The calcium indicators were excited by light from a 100 W mercury short arc lamp and the resulting fluorescence was measured by a charge-coupled device (CCD) camera at a frame rate of 5 Hz (Rolera XR, Qimaging, Surrey, BC, Canada). We used a 16 X/0.8 NA water-immersion objective lens for imaging (CFI75 LWD 16XW; Nikon Instruments). Three to five trials were averaged for each visual stimulus.

Scopolamine, resp. muscarine were prepared at 6 mM in locust saline containing fast green (0.5%) to visually monitor the affected region. They were puffed using a pneumatic picopump (PV830, WPI, Sarasota, FL). Injection pipettes had tip diameters of approximately 2 µm and were visually positioned between the excitatory dendritic field of the LGMD and its most superficial inhibitory field with a micromanipulator (fields A and C; O’Shea and Williams, 1974). Brief air pulses with a pressure of ~6 psi (41.4 kPa) caused drug ejection into the optic lobe. Spread of fast green within the lobe was monitored to ensure drugs diffused through the excitatory dendritic field. Based on previous calculations of drug dilution within the optic lobe (see Methods in Dewell and Gabbiani, 2017), our best estimate for the final drug concentration at the LGMD is ~100 µM.

Intracellular LGMD and extracellular DCMD recordings were performed simultaneously with calcium imaging in some animals. After identification of the location in the LGMD’s excitatory dendritic branches that had been stained with the calcium indicator OGB-5N, a sharp electrode (25 MΩ) containing 3 µl of 2 M potassium acetate, 0.5 M potassium chloride and 0.1 mM Alexa 488 was inserted into the excitatory dendrite of the LGMD under the 16 X objective lens for recording. Intracellular signals were amplified in bridge mode with an Axoclamp 2B and an instrumental amplifier (Brownlee Precision 440; NeuroPhase, Palo Alto, CA). LGMD and DCMD signals were acquired with a sampling rate of 5 kHz by a data acquisition card (PCI 6110; National Instruments, Austin, TX).

### Data analysis

Data analysis was performed with Matlab. The relative fluorescence change was calculated as dF/F(t)=[F(t)-F_0_]/F_0_, with the baseline fluorescence F_0,_ being the average of the first 10 frames (2 s) before the visual stimulus. For peak dF/F spatial maps (e.g. Fig. 2A), the value of dF/F(t) at each pixel was calculated as the median over a 5 × 5-pixel area centered at the given pixel (0.9 µm/pixel). After median filtering, only suprathreshold pixels were used, defined as having a peak dF/F > 180% of the maximum noise of dF/F at that pixel. The maximum noise at each pixel was computed as the maximum of the dF/F values within 2 s before the onset of the visual stimulus. The threshold value of 180% was selected to eliminate pixels resulting in falsely positive dF/F outside the dendritic branches based on earlier, two-photon imaging experiments with high spatial resolution (Zhu and Gabbiani 2016, Fig. 1D). The peak dF/F values at the pixels above threshold were retained and passed through a Gaussian filter with a standard deviation sigma = 5 pixels. In Fig. 1D, we selected the threshold value (dashed line) to be well above noise level, so that calcium signals were unambiguously increasing at those time points. The results in Fig. 1E and F are robust to changes of the threshold around that value.

To map the dendritic receptive fields of square flashes at 5 locations on the screen, we first measured the peak dF/F spatial map in response to individual square flashes. For this purpose, we first filtered the pixels that were above the noise level (see above), then computed the maximum of the peak dF/F over all the pixels and retained those pixels above 4% of the maximum. This additional thresholding step was necessary due to the small amplitude of peak dF/F in response to square flashes. The threshold was selected to eliminate falsely positive dF/F pixels outside the dendritic branches. The branches that were covered by the retained pixels were taken to be the dendritic receptive fields of that square flash.

The principal axes of the peak dF/F spatial map were computed by first scaling and rounding the intensity of the peak dF/F at each pixel into integers. Next, we took these integers for each pixel as the number of identical data points with coordinates matching those of that pixel. After that, we performed principal component analysis on all these data points to extract the first principal component that had the largest variance and the second principal component orthogonal to the first one. We rotated the peak dF/F spatial map to make the first or second principal axis horizontal, and summed the values of peak dF/F along the vertical axis to get the projected values along each principal axis.

To make plots, the membrane potential was first median-filtered over a time window of 30 ms and down-sampled 100 times (to 50 Hz).

### Statistics

For paired comparison of the control and scopolamine groups, we performed a nonparametric one-tailed Wilcoxon sign-rank test. The null hypothesis was rejected and the results were judged statistically significant at probabilities p<0.05. For responses to small flashes (Supp. Fig. 3G-I) a non-parametric Wilcoxon rank sum test was conducted on the number of spikes in response to individual stimuli before and after addition of scopolamine. For determining whether there are any statistically significant differences between the means of groups stimulated with gratings of varying separations, we performed a one-way analysis of variance (one-way ANOVA). The null hypothesis that the means of all groups were statistically equal was rejected at significance levels p<0.05. For comparing the slopes of two regression lines, we performed an analysis of covariance (ANCOVA). The null hypothesis that the slopes of the two regression lines were statistically equal was rejected at significance levels p<0.05. A two-way analysis of variance (two-way ANOVA) was performed to determine whether there were any statistically significant differences between the control and the scopolamine/muscarine groups, each of which having different levels (types of visual stimuli). A value of p<0.05 was regarded as significant. For experiments studying the influence of muscarine on calcium responses to flicker-off stimuli with different sizes, statistical analysis of paired comparisons for each stimulus type was not performed as the separation between control and muscarine was unambiguous already after testing 3 animals (Fig. 6E, F). For comparisons using the paired t-test, normality was tested with a Lilliefors test with an alpha level of 0.2. Throughout, we use the abbreviation sd for standard deviation.

## Acknowledgements

We thank Dr. E. Froudarakis for comments on the manuscript. The work was supported by grants from the National Science Foundation (DMS-1120952), the National Institutes of Health (NIH) (MH-065339) and NEI Core Grant for Vision Research (EY-002520-37).

## Author Contributions

Y.Z., R.B.D. and F.G. conceived the experiments; Y.Z. performed calcium imaging experiments; Y.Z., R.B.D and H.W. performed electrophysiology experiments; Y.Z. and R.B.D. analyzed data; Y.Z., R.B.D. and F.G. prepared figures and wrote the paper.

## Competing interests

No competing interests declared.

